# Medial prefrontal cortex stimulation abolishes implicit reactions to threats and prevents the return of fear

**DOI:** 10.1101/2023.02.06.527256

**Authors:** Eugenio Manassero, Giulia Concina, Maria Clarissa Chantal Caraig, Pietro Sarasso, Adriana Salatino, Raffaella Ricci, Benedetto Sacchetti

**Affiliations:** Rita Levi-Montalcini Department of Neuroscience, University of Turin, Corso Raffaello 30, 10125 Turin, Italy; Department of Psychology, University of Turin, Via Giuseppe Verdi 10, 10124 Turin, Italy

## Abstract

Down-regulating emotional overreactions toward threats is fundamental for developing treatments for anxiety and post-traumatic disorders. The prefrontal cortex (PFC) is critical for top-down modulatory processes, and despite previous studies adopting repetitive Transcranial Magnetic Stimulation (rTMS) over this region provided encouraging results in enhancing extinction, no studies have hitherto explored the effects of stimulating the medial PFC (mPFC) on threat memory and generalization. Here we showed that rTMS applied before threat memory retrieval abolishes implicit reactions to learned and novel stimuli in humans. These effects were not due to inhibition of electrodermal reactivity and enduringly persisted one week later in the absence of rTMS. No effects were detected on explicit recognition. Critically, we observed stronger attenuation of defensive responses in subjects stimulated over the mPFC than the dlPFC. Our findings uncover a prefrontal region whose modulation can permanently hamper implicit reactions to learned dangers, representing an advance to long-term deactivating overreactions to threats.

## Introduction

Emotional memories related to past threat experiences allow humans to predict future dangers and trigger adaptive defensive reactions when encountering learned threat-signaling cues^1^. However, extremely dangerous situations may represent points of origin for trauma and lead to maladaptive behaviors and psychological disorders^2^. In this scenario, intrusive traumatic memories may be explained by over-time enduring conditioned responses to trauma reminders^3^. Attempting to down-regulate the emotional overreactions toward threat-predictive stimuli is one of the main routes for developing effective treatments for anxiety and post-traumatic disorders. Common approaches such as pharmacological treatments and cognitive-behavioral therapy (CBT) have demonstrated partial efficacy^4^, and recent evidence suggests that the functional outcome of behavioral methods may depend on the extent to which the prefrontal cortex is recruited during these processes^5^. Hence, new intervention strategies influencing the prefrontal dynamics would represent an important advance in the field. One recently argued possibility consists of modulating prefrontal and other neural networks underlying threat memory processes through non-invasive brain stimulation^6^.

Previous studies adopted transcranial direct current stimulation (tDCS) or transcranial electrical stimulation (tES) to disrupt the consolidation of these memories^7‒9^, potentiate extinction processes^10, 11^, and narrow threat generalization patterns^12^, leading to contradictory results. According to one work^7^, cathodal stimulation over the dorsolateral prefrontal cortex (dlPFC) disrupted threat memory consolidation, with no enhancing effect of anodal stimulation. In contrast, other studies found an increase in implicit responses with anodal stimulation^8^ and no effect of cathodal stimulation^9^ over the same site. Moreover, one study employing anodal stimulation over the dlPFC^11^ revealed an improvement in extinction learning but no delayed effects on the recall of the extinction memory. A further investigation^10^ reported detrimental effects of stimulating the medial prefrontal cortex (mPFC), i.e. low-frequency alternating-current (AC) stimulation augmented the defensive responses, whereas direct-current (DC) stimulation widened threat generalization profiles.

An alternative neurostimulation approach is repetitive transcranial magnetic stimulation (rTMS), which ensures greater focality^13, 14^. Some rTMS studies targeted the mPFC^15^ and the posterior PFC^16^ to obtain a successful enhancement of extinction learning, while others^17^ targeted the dlPFC to disrupt threat-memory reconsolidation. Indeed, most rTMS-based research targeting the PFC has pursued an improvement of fear extinction and no previous studies attempted to down-modulating the defensive responses triggered by a learned threatening stimulus without adopting fear extinction.

So far, human brain stimulation studies have been mainly focused on the dorsolateral region of the PFC^6^. However, within the PFC, a brain structure that has emerged to be critically engaged in the downstream regulation of subcortical threat detection systems (such as amygdalar nuclei) is the medial prefrontal cortex (mPFC)^18^. The mPFC is a neural region that has received enormous attention in the social, cognitive, and emotional domains of neuroscientific research^19^. Several studies highlighted its role in threat extinction^20‒22^, threat generalization^23^, threat learning^24^, and both rodent and human studies^25, but see^ ^6^ suggest that the neural enhancement of the mPFC has a strong potential to augment extinction and reduce fear. However, to our knowledge, no study has been so far conducted to explore the effects of mPFC stimulation on the expression of a threat memory without extinction learning in humans.

To explore this possibility, here we tested whether an rTMS protocol applied over the mPFC may enduringly attenuate the defensive responses to a learned threat-predictive stimulus.

## Results

### mPFC-focused rTMS effects on implicit reactions toward threat-predictive cues

To explore the effects of an mPFC-centered rTMS on the defensive responses to a learned threat, we designed a three-session experiment starting with a threat learning followed by an implicit retention test and a follow-up implicit re-test (Figure 1).

**Figure 1.**
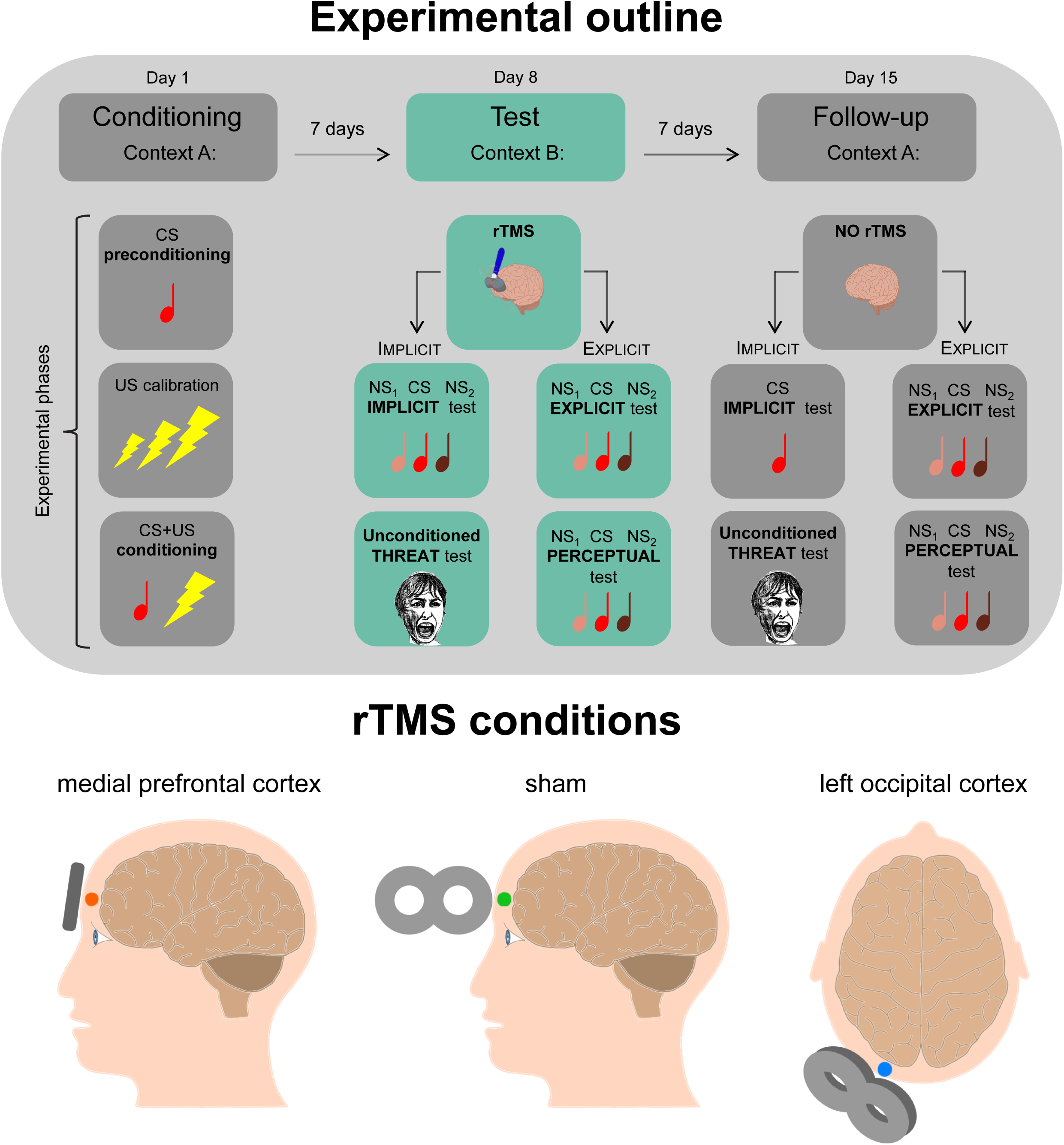
Schematic diagram depicting the experimental outline and rTMS conditions. In the first session (day 1, context A), participants underwent a single-cue threat conditioning in which a tone (CS) was paired with a mild electrical shock (US). In the second session (day 8, context B), a 1Hz-rTMS procedure was actively applied over the medial prefrontal cortex (mPFC, *n* = 21; mPFC-E, *n* = 21), sham-applied over the same site (sham, *n* = 21; sham-E, *n* = 21), or actively applied to the left occipital cortex (OC, *n* = 21). In the implicit conditions (mPFC, sham, OC), subjects underwent an implicit test during which they were presented with the CS and two new stimuli (NS_1_ and NS_2_) and then an unconditioned threat test while being recorded in their SCRs. In the explicit conditions (mPFC-E, sham-E), participants underwent an explicit 2AFC recognition task during which they were presented with tone pairs each composed of the CS and one of the two NSs, and they were asked to recognize the CS providing a confidence level for each choice. Last, participants underwent a 2AFC perceptual discrimination test, in which they had to judge whether the two tones in each pair (CS and/or NSs) were “the same tone” or “different tones”. The third session (day 15, context A) was identical to the second one except for the absence of the rTMS.

During the conditioning session, participants learned to associate an auditory cue (conditioned stimulus, CS, 800Hz) with a mild electric stimulation (unconditioned stimulus, US, individually calibrated intensity) in a given environment (context A). We adopted a single-cue conditioning paradigm because it more ecologically reflects real-life traumatic experiences^26‒30^. To validate the between-groups homogeneity in the painful stimuli perception, we computed a one-way ANOVA on post-conditioning US ratings, and we observed no significant differences amongst groups (F_(2,60)_ = 0.6108, *P* > 0.05, η_p_^2^ = 0.01995) (Table 1). We also did not observe significant differences amongst groups in SCRs to the CS during the preconditioning phase (F_(2,60)_ = 1.539, *P* > 0.05, η_p_^2^ = 0.04879) nor to the US during the conditioning phase (F_(2,60)_ = 1.237, *P* > 0.05, η_p_^2^ = 0.03961) (Supplementary Figure 1).

**Table 1.** Experimental groups’ descriptive, experimental and clinical data. The table reports, for each experimental condition: sample size (*N*), sex distribution (F = Female, M = Male), mean age, State-Trait Anxiety Inventory Form Y (STAI-Y) State subscale score during session 1 (S1), session 2 (S2), and session 3 (S3), and Trait subscale score, US current intensity (mA), post-conditioning US rating, TMS resting motor threshold (rMT), and TMS power. All data are mean ± standard deviation.

| Group | *N* | Sex | Age | STAI-Y<br>State<br>(S1) | STAI-Y<br>State<br>(S2) | STAI-Y<br>State<br>(S3) | STAI-Y<br>Trait | US<br>(mA) | US<br>rating | TMS<br>rMT | TMS<br>power |
| --- | --- | --- | --- | --- | --- | --- | --- | --- | --- | --- | --- |
| mPFC | 21 | 13F<br>8M | 22.93<br>± 2.09 | 30.29<br>± 3.51 | 32.24<br>± 7.23 | 30.38<br>± 6.41 | 39.10<br>± 6.71 | 4.94<br>± 2.24 | 5.29<br>± 0.94 | 56.38<br>± 6.30 | 39.81<br>± 0.51 |
| sham | 21 | 12F<br>9M | 22.96<br>± 2.34 | 33.86<br>± 5.44 | 32.29<br>± 5.92 | 31.90<br>± 6.61 | 39.14<br>± 3.80 | 5.20<br>± 2.64 | 5.52<br>± 0.86 | - | - |
| OC | 21 | 13F<br>8M | 24.00<br>± 2.82 | 31.14<br>± 4.71 | 31.86<br>± 8.52 | 31.29<br>± 6.96 | 39.10<br>± 5.49 | 4.47<br>± 3.28 | 5.24<br>± 0.89 | 61.05<br>± 4.66 | 40.00<br>± 0.00 |
| mPFC-E | 21 | 13F<br>8M | 24.39<br>± 2.43 | 31.71<br>± 4.89 | 30.90<br>± 5.66 | 30.48<br>± 4.96 | 38.29<br>± 6.21 | 5.13<br>± 1.86 | 5.43<br>± 0.94 | 58.67<br>± 7.16 | 39.52<br>± 1.54 |
| sham-E | 21 | 13F<br>8M | 23.83<br>± 2.73 | 33.10<br>± 5.59 | 31.48<br>± 5.54 | 30.38<br>± 7.73 | 38.29<br>± 5.22 | 5.27<br>± 3.19 | 5.31<br>± 1.31 | - | - |
| dIPFC | 21 | 14F<br>7M | 23.90<br>± 3.14 | 31.52<br>± 4.38 | 30.38<br>± 6.36 | 30.71<br>± 5.72 | 39.05<br>± 5.95 | 4.36<br>± 1.59 | 5.55<br>± 1.21 | 57.71<br>± 5.73 | 39.86<br>± 0.48 |

One week later, we tested the implicit memory of the learned association in control subjects and in those that received rTMS over the mPFC. To locate this brain region, which corresponds to the Brodmann area 10^31^, we positioned the coil over Fpz adopting the 10‒20 EEG coordinate system, since previous rTMS studies^15, 32, 33^ ensured this placement reached the mPFC. An offline 10-min session of 1Hz-rTMS targeting this neural site (mPFC, *n* = 21) was applied immediately before memory retrieval. Besides proximal inhibitory effects on local brain activity, 1Hz-rTMS protocols have been shown to induce downstream distal effects through the potentiation of resting-state functional connectivity with remote brain areas inside the stimulated neural network^34^. When applied over prefrontal regions, 1Hz-rTMS seems to increase regional cerebral blood flow (rCBF)^35, 36^. Control subjects underwent a 10-min sham stimulation procedure over the same cortical area (sham, *n* = 21). To ascertain the topographical selectivity, in one further condition (OC, *n* = 21) we applied the rTMS over the left occipital cortex as an active control site.

Memory retention was tested in a different environment from that where the learning had occurred (context B) to avoid any contextual influence on retrieval^37‒41^. Indeed, the context shift for this session mirrors a real-life treatment setting ‒which unlikely takes place in the threatening location. To test implicit threat memory, we performed an implicit recognition task in which subjects were exposed to the CS while being recorded in their evoked autonomic reactions (i.e., electrodermal skin conductance responses, SCRs). No US shocks were delivered during this phase. Besides the CS, participants were presented with two novel but perceptually similar tones (NS_1_, 1000Hz; NS_2_, 600Hz) to study threat generalization. Auditory frequencies of NSs were selected to obtain a slowly decaying gradient of defensive tunings^39, 42, 43^. A one-way ANOVA (F_(2,60)_ = 8.793, *P* < 0.001, η_p_^2^ = 0.2267) revealed that subjects that received rTMS over the mPFC exhibited weakened SCRs than those observed in the sham (*P* < 0.001) and OC (*P* = 0.0074) groups, where the CS evoked similarly strong autonomous reactions (*P* > 0.05) (Figure 2A). These data were obtained by averaging all four CS trials. To test potential differences in the genesis of each CS-related response, we performed a trial-by-trial analysis through a 3 × 4 mixed ANOVA. This analysis revealed a significant main effect of group (F_(2,60)_ = 8.793, *P* < 0.001, η_p_^2^ = 0.2267), a significant main effect of trial (F_(3,180)_ = 12.328, *P* < 0.001, η_p_^2^ = 0.17) and a not significant group × trial interaction (F_(6,180)_ = 1.155, *P* > 0.05, η_p_^2^ = 0.037). Notably, we observed that in Trial 1 the mPFC group already exhibited weaker reactions than sham (*P* = 0.002) and OC (*P* = 0.046) conditions, which did not differ from each other (*P* > 0.05) (Figure 2B). This data indicates that the rTMS procedure immediately affected SCRs triggered by memory retrieval, rather than interfering with reactivation-induced reconsolidation processes, as previously reported^17^. To the best of our knowledge, this is the first evidence that brain stimulation may promptly attenuate implicit defensive reactions during memory retrieval, without requiring repeated exposure to the CS (i.e., extinction learning).

**Figure 2.**
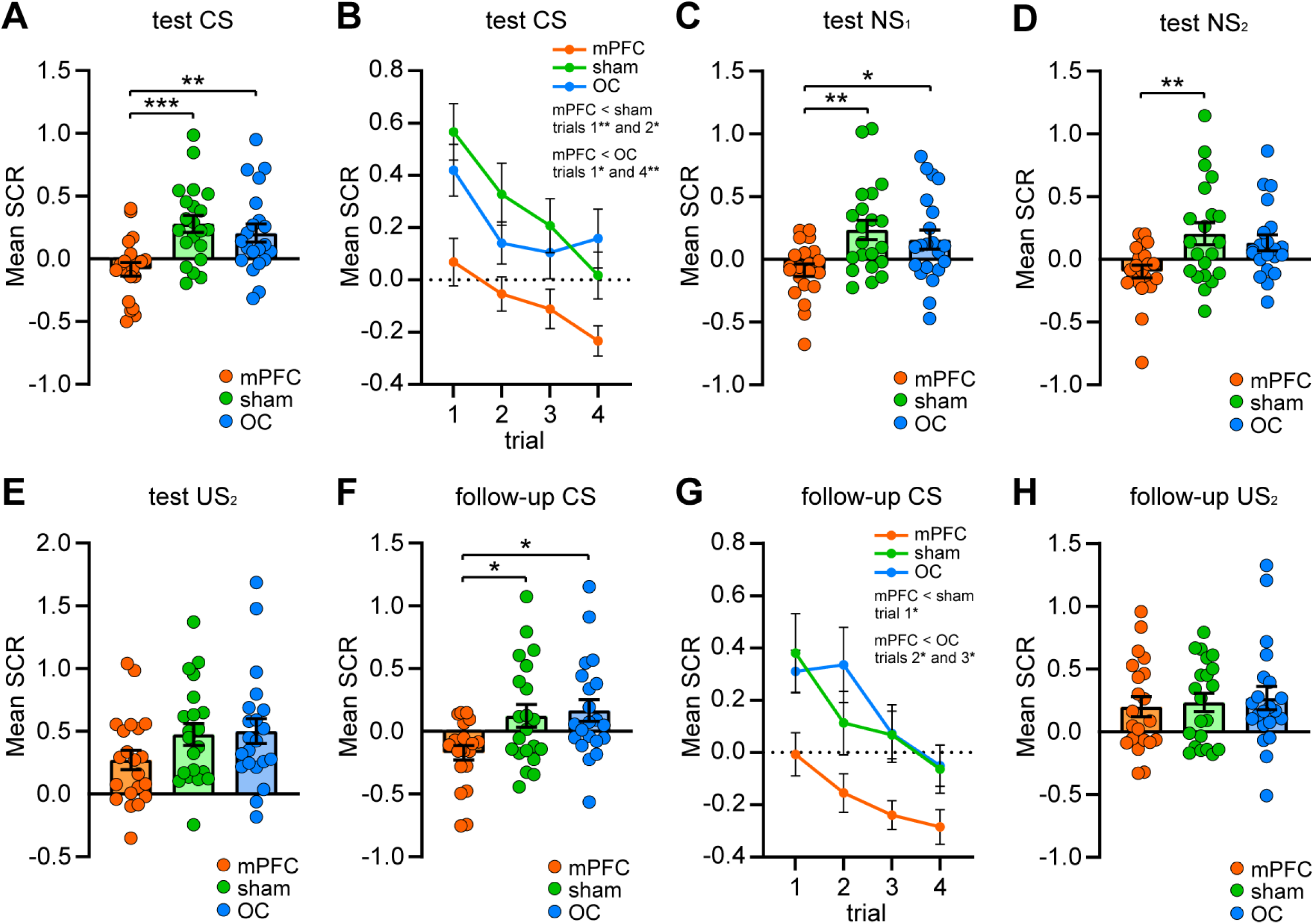
Effects of rTMS over the mPFC on immediate and remote implicit threat memory, threat generalization to new stimuli, and overall electrodermal responsivity. (**A**) Dot plot representing the mean SCRs elicited by the CS during the test session in the three different conditions. The group stimulated over the mPFC showed attenuated implicit reactions relative to both sham and OC conditions. (**B**) Time-course of CS trials (1‒4) during the test session in the three different groups. Already in the first presentation of the CS, the mPFC group displayed dampened responses relative to sham and OC groups. (**C**) Implicit reactions to the NS_1_ during the test session were reduced in the mPFC group relative to the sham and OC groups. (**D**) Implicit reactions to the NS_2_ during the test session were lower in the mPFC group than in the sham group. (**E**) Implicit reactions to the US_2_ during the test session were similar among conditions, showing no immediate rTMS effects on the overall electrodermal responsivity. (**F**) In the follow-up session, the mPFC group enduringly demonstrated attenuated implicit reactions to the CS. (**G**) Time-course of CS trials (1‒4) during the follow-up session in the three different groups. (**H**) Implicit reactions to the US_2_ during the follow-up session were comparable among groups. * *P* < 0.05, ** *P* < 0.01, *** *P* < 0.001. All data are mean and SEM. In each graph, the level “0” corresponds to the pre-conditioning mean SCR response to the CS. One-way ANOVA followed by Bonferroni-adjusted *post hoc* comparisons (A, C, D, E, F, H); 3×4 mixed ANOVA followed by Bonferroni-adjusted *post hoc* comparisons (B, G).

In the same session, we also analyzed threat generalization to the NSs, and we found that the mPFC group displayed attenuated responses to the NS_1_ relative to the other two conditions, which did not differ between each other (F_(2,60)_ = 5.856, *P* = 0.0048, η_p_^2^ = 0.1633; mPFC vs sham: *P* = 0.0051; mPFC vs OC: *P* = 0.0483; sham vs OC: *P* > 0.05) (Figure 2C). Autonomous responses to the NS_2_ (F_(2,60)_ = 5.20, *P* = 0.0083, η_p_^2^ = 0.1477) were reduced in the mPFC group relative to the sham group (*P* = 0.0091) but not relative to the OC group (*P* = 0.067), whereas sham and OC groups were similar (*P* > 0.05) (Figure 2D). We then explored fear tunings (i.e. the decay of defensive responses typically observed by increasing a cue’s perceptual distance from the CS, where the decay rate reflects the strength of threat generalization)^42^ by comparing within-subjects response amplitudes to the learned and new tones. We found that defensive responses were overall reduced (repeated measures ANOVA, F_(1.734,34.68)_ = 0.07933, *P* > 0.05, η_p_^2^ = 0.003951) in the mPFC group since SCRs evoked by the CS were similarly low to those triggered by the NS_1_ and the NS_2_. The tuning strength was spread to both NSs in the sham (F_(2,40)_ = 0.6713, *P* > 0.05, η_p_^2^ = 0.03248) as well as in the OC condition (F_(2,40)_ = 1.777, *P* > 0.05, η_p_^2^ = 0.08162), thus indicating threat generalization. These data point to an attenuating effect of the pre-retrieval mPFC-focused rTMS on the immediate implicit defensive responses to the learned threatening cue as well as to new perceptually similar stimuli.

We next sought to disambiguate whether the rTMS effects were due to a general down-regulation of electrodermal responsivity, or whether they specifically targeted the threat memory. To this end, subjects were presented with an unconditioned threatening stimulus consisting of a female scream sample (unconditioned stimulus 2, US_2_) while being recorded in their SCRs. A one-way ANOVA revealed no differences amongst conditions (F_(2,60)_ = 2.045, *P* > 0.05, η_p_^2^ = 0.06381), indicating that the rTMS did not cause an overall inhibition of electrodermal reactivity (Figure 2E).

### rTMS long-lasting effects on implicit threat memory

To test whether and to what extent rTMS-related outcomes endured beyond the short-term after-effect window and persisted in a long-term period, we planned a follow-up session. One week after rTMS session and threat memory retrieval test, all participants returned to the conditioning room (context A) and underwent a re-testing phase, which was identical to the testing one except for the absence of rTMS administration. This phase allowed us to evaluate potential long-term rTMS effects, and to test a possible renewal effect since other approaches based on extinction learning are context-dependent^44^ and here subjects were re-exposed to the original threatening environment.

Concerning the implicit responses to the CS, participants of the mPFC group persisted in displaying weaker SCRs (F_(2,60)_ = 5.421, *P* = 0.0069, η_p_^2^ = 0.153) than those observed in the sham (*P* = 0.0315) and OC (*P* = 0.0110) groups. On the contrary, the CS elicited comparable autonomous reactions in the sham and the OC conditions (*P* > 0.05) (Figure 2F). Trial-by-trial analysis through the 3 × 4 mixed ANOVA revealed a significant main effect of group (F_(2,60)_ = 5.421, *P* = 0.0069, η_p_^2^ = 0.153), a significant main effect of trial (F_(3,180)_ = 14.915, *P* < 0.001, η_p_^2^ = 0.199) and a not significant group × trial interaction (F_(6,180)_ = 1.09, *P* > 0.05, η_p_^2^ = 0.035) (Figure 2G). We next analyzed the autonomous response patterns to the US_2_ and again we found comparable reactions amongst the groups (F_(2,60)_ = 0.1787, *P* > 0.05, η_p_^2^ = 0.005921) (Figure 2H). These findings support an enduring effect of the mPFC-rTMS in attenuating the long-term implicit defensive responses to the learned threat-predictive cue, independently from the context-shift for rTMS administration and even with the re-exposition to the environment where threat learning had occurred. This persistent effect was expressed notwithstanding an unaffected electrodermal overall reactivity.

### mPFC-focused rTMS effects on explicit memory recognition and perceptual discrimination of threat-predictive cues

We then investigated the effect of rTMS over the mPFC on the retention of explicit-declarative threat memories. A further group of subjects that received the identical 1Hz-rTMS procedure over the mPFC (mPFC-E, *n* = 21) and a further control group (sham-E, *n* = 21) (which had reported similar post-conditioning US ratings: *t*_(40)_ = 0.3387, *P* > 0.05, η_p_^2^ = 0.00286) immediately underwent an explicit two-alternative forced-choice (2AFC) recognition task, in which they were presented with a random sequence of tone pairs, each composed of the CS and one of the two NSs. Here, subjects were asked to consciously identify and refer which stimulus of each pair was the one previously paired with the US (i.e., the CS), and to provide a subjective confidence level for each choice using a scale ranging from 0 (completely unsure) to 10 (completely sure)^39, 45^. Explicit recognition patterns revealed that each experimental condition successfully identified the CS amongst the NSs with an accuracy level above the 50% chance level (mPFC-E: *t*_(20)_ = 9.226, *P* < 0.0001, η_p_^2^ = 0.8098; sham-E: *t*_(20)_ = 14.24, *P* < 0.0001, η_p_^2^ = 0.9103). A between-groups comparison (*t*_(40)_ = 1.114, *P* > 0.05, η_p_^2^ = 0.03011) showed no differences in the explicit recognition accuracy (Figure 3A). Confidence ratings provided by participants supported the lack of rTMS-related effects since the two groups were similarly confident when making their choices (*t*_(40)_ = 0.8419, *P* > 0.05, η_p_^2^ = 0.01741) (Figure 3B).

**Figure 3.**
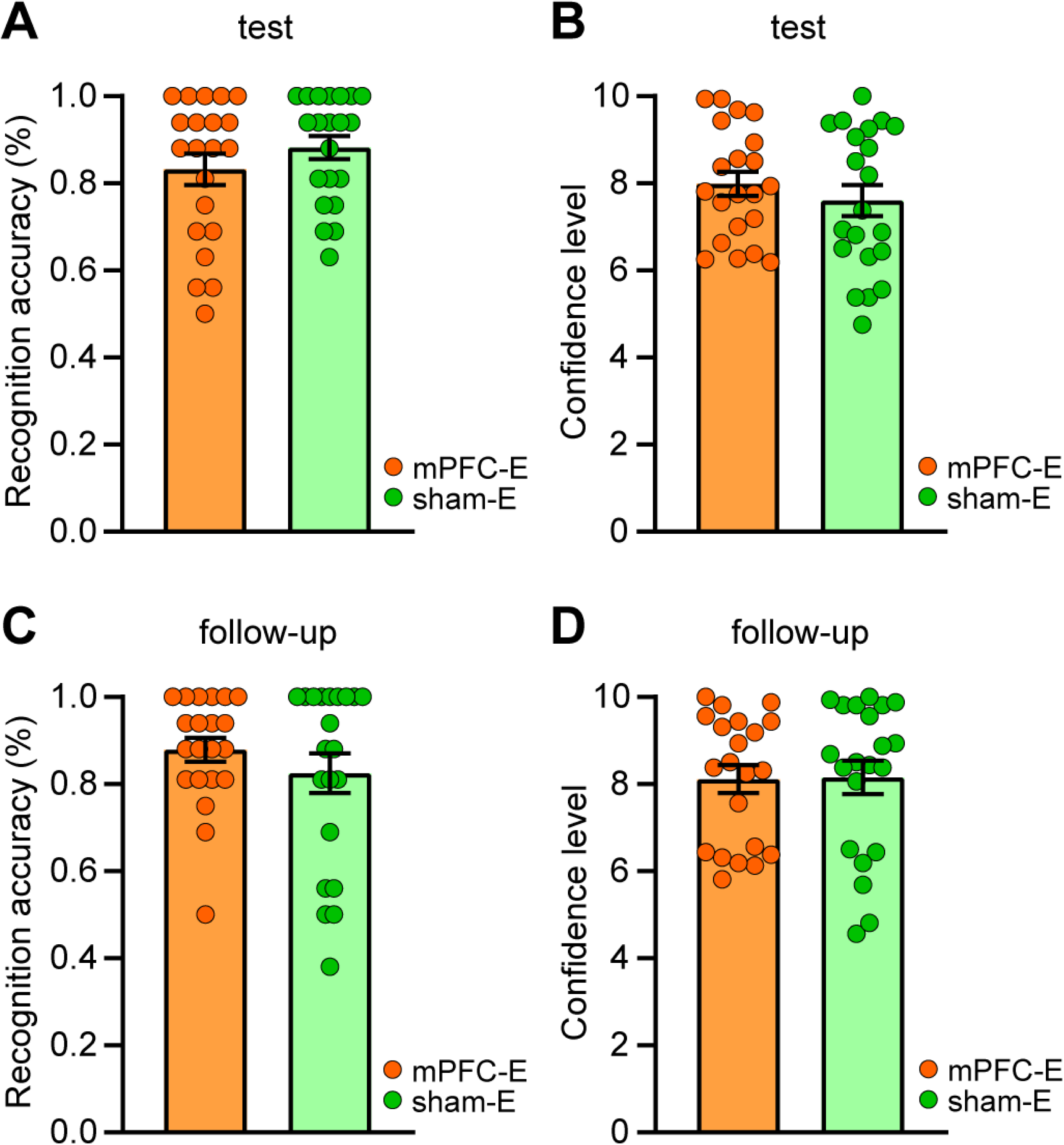
Effects of rTMS over the mPFC on immediate and remote explicit threat memory. (**A**) During the test session, explicit recognition patterns were comparable between the group stimulated over the mPFC and the sham group. (**B**) During the test session, confidence ratings were similar between the two conditions. (**C**) During the follow-up session, mPFC-E and sham-E groups similarly identified the CS between the NSs. (**D**) During the follow-up session, mPFC-E and sham-E groups were comparably confident about their explicit choices. All data are mean and SEM. Student’s unpaired *t* test (A, B, C, D).

Next, we implemented a 2AFC perceptual task in which we investigated the ability of participants to sensory discriminate between the CS and the two NSs by collecting binary ‘same or different’ judgments as well as confidence ratings. The perceptual discrimination test yielded no significant between-groups differences in accuracy (*t*_(40)_ = 1.362, *P* > 0.05, η_p_^2^ = 0.04434) as well as confidence levels (*t*_(40)_ = 0.9165, *P* > 0.05, η_p_^2^ = 0.02057). Indeed, both groups discriminated the CS from the NSs with high precision (mPFC-E: 0.9795 ± 0.01503 SEM; sham-E: 1.00 ± 0.00 SEM) and with comparably high confidence levels (mPFC-E: 9.409 ± 0.1529 SEM; sham-E: 9.586 ± 0.1173 SEM), thereby showing no rTMS effects on sensory abilities.

These data suggest that the pre-retrieval rTMS procedure over the mPFC did not affect the explicit recognition nor the perceptual discrimination of a learned threat. This dissociation is in line with the idea that implicit and explicit threat memory systems are mediated by distinct neural circuits^46‒48^ which may lead to divergent defensive responses even in presence of new stimuli^39^.

During the follow-up session, explicit recognition patterns demonstrated an over-chance accuracy level for each group (mPFC-E: *t*_(20)_ = 13.78, *P* < 0.0001, η_p_^2^ = 0.9047; sham-E: *t*_(20)_ = 7.162, *P* < 0.0001, η_p_^2^ = 0.7195). Again, here there were no between-groups differences (*t*_(40)_ = 1.024, *P* > 0.5, η_p_^2^ = 0.02553) since both groups achieved a comparably high recognition accuracy (Figure 3C). Each group also reported similar confidence levels (*t*_(40)_ = 0.08394, *P* > 0.05, η_p_^2^ = 0.0001761) (Figure 3D).

As in the case of the previous session, we did not observe significant between-group differences in the perceptual discrimination (*t*_(40)_ = 1.00, *P* > 0.05, η_p_ = 0.02439) and the respective confidence ratings (*t*_(40)_ = 0.1489, *P* > 0.05, η_p_ = 0.0005542). Indeed, the discrimination accuracy (mPFC-E: 1.00 ± 0.00 SEM; sham-E: 0.9933 ± 0.006667 SEM) and the self-assessed confidence (mPFC-E: 9.598 ± 0.1466 SEM; sham-E: 9.633 ± 0.1816 SEM) were similarly high in each condition.

These findings suggest that the rTMS procedure over the mPFC did not affect the long-term capacity to consciously identify and perceptually discriminate the learned threat-predictive cue.

### Effects of rTMS over medial and dorsolateral prefrontal cortex on immediate and long-term implicit expression of a threat memory

Next, we asked whether the findings we obtained by targeting the mPFC were finely specific for this site or, alternatively, they overlapped with those observed by targeting other prefrontal sub-regions. For this purpose, in one further group (dlPFC, *n* = 21) we applied the same rTMS procedure over the left dorsolateral PFC (Figure 4A) and we then compared the implicit patterns of this group with those displayed by the mPFC condition. We selected the left dlPFC since previous studies^e.g. 16^ targeted the left hemisphere for testing the rTMS effects on the PFC, and some evidence^see^ ^6^ suggested that inhibitory tDCS and rTMS over the left dlPFC may disrupt threat memory consolidation.

**Figure 4.**
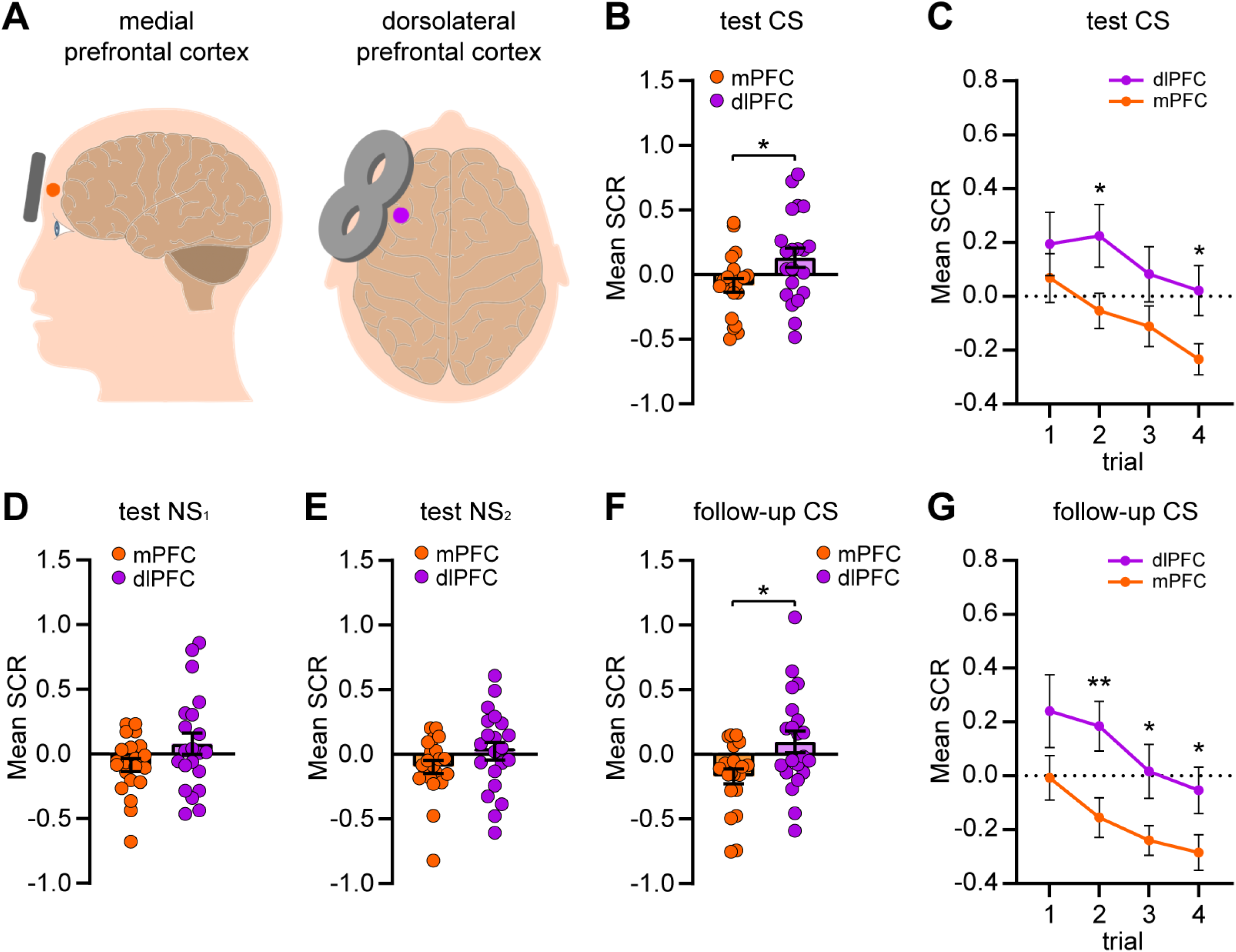
Different effects of rTMS over the mPFC and the left dlPFC on immediate and remote implicit threat memory. (**A**) One week after threat learning (day 1, context A), a 1 Hz rTMS procedure was actively applied over the left dorsolateral prefrontal cortex (dlPFC, *n* = 21) and compared with the same mPFC-stimulated group of Fig. 2 (mPFC, *n* = 21) (day 8, context B). Both groups underwent the implicit test and, one further week later, the follow-up implicit test (day 15, context A). (**B**) Dot plot representing the mean SCRs elicited by the CS during the test session in the two different conditions. The group stimulated over the mPFC showed weaker implicit reactions than the group stimulated over the dlPFC. (**C**) Time-course of CS trials (1‒4) during the test session in the two different groups. (**D, E**) Implicit reactions to the NS_1_ (**D**) and the NS_2_ (**E**) during the test session were comparable in the two groups. (**F**) During the follow-up session, the group stimulated over the mPFC persisted to show weaker implicit responses to the CS than the group stimulated over the dlPFC. (**G**) Time-course of CS trials (1‒4) during the follow-up session in the two different groups. * *P* < 0.05, ** *P* < 0.01. All data are mean and SEM. In each graph, the level “0” corresponds to the pre-conditioning mean SCR response to the CS. Student’s unpaired *t* test (B, D, E, F); 2×4 mixed ANOVA followed by Bonferroni-adjusted *post hoc* comparisons (C, G).

First, we find no significant differences between the two conditions in the post-conditioning US ratings (*t*_(40)_ = 0.7809, *P* > 0.05, η_p_^2^ = 0.01502) (Table 1), in SCRs to the CS during the preconditioning phase (*t*_(40)_ = 0.03252, *P* > 0.05, η_p_^2^ = 0.00002643) and to the US during the conditioning phase (*t*_(40)_ = 1.905, *P* > 0.05, η_p_^2^ = 0.0832) (Supplementary Figure 2). Then, we compared the implicit reactions toward the CS during the test session, and we found weaker defensive responses in the mPFC group relative to the dlPFC group (*t*_(40)_ = 2.319, *P* = 0.0256, η_p_^2^ = 0.1185) (Figure 4B). A 2 × 4 mixed ANOVA model on the genesis of each CS-evoked response revealed a significant main effect of group (F_(1,40)_ = 5.376, *P* = 0.026, η_p_^2^ = 0.118), a significant main effect of trial (F_(2.41,96.389)_ = 3.921, *P* = 0.017, η_p_^2^ = 0.089), and a not significant group × trial interaction (F_(2.41,96.389)_ = 0.402, *P* > 0.05, η_p_^2^ = 0.010) (Figure 4C).

We found no between-groups differences in the implicit responses to the NS_1_ (*t*_(40)_ = 1.72, *P* > 0.05, η_p_^2^ = 0.06883) (Figure 4D) and the NS_2_ (*t*_(40)_ = 1.416, *P* > 0.05, η_p_^2^ = 0.04774) (Figure 4E), showing that the divergent rTMS effects in the mPFC and the dlPFC groups were selective for the CS. Fear tuning analysis of dlPFC group’s implicit reactions revealed similar SCRs amplitudes elicited by the CS and both NSs (F_(2,40)_ = 2.856, *P* > 0.05, η_p_^2^ = 0.1249).

This different pattern toward the learned threatening cue was replicated during the follow-up session since the mPFC group persisted in more dimly reacting to the CS with respect to the dlPFC group (*t*_(40)_ = 2.650, *P* = 0.0115, η_p_^2^ = 0.1493) (Figure 4F). Trial-by-trial analysis through the 2 × 4 mixed ANOVA revealed a significant main effect of group (F_(1,40)_ = 7.023, *P* = 0.011, η_p_^2^ = 0.149), a significant main effect of trial (F_(1.957,78.266)_ = 8.708, *P* < 0.001, η_p_ = 0.179) and a not significant group × trial interaction (F_(1.957,78.266)_ = 0.310, *P* > 0.05, η_p_^2^ = 0.008) (Figure 4G). Thus, these findings demonstrated that without adopting an extinction paradigm as in other previous studies^15‒17^, there was a more pronounced and enduring attenuation of implicit defensive reactions to a threat-predictive stimulus when the rTMS was delivered over the medial than over the left dorsolateral PFC.

## Discussion

In this study, we found that implicit reactions to both learned and novel stimuli were reduced following a 1Hz-rTMS procedure over the mPFC.

So far, most rTMS studies targeting this prefrontal area have been conducted to enhance fear extinction processes. Indeed, a large body of findings suggests that the prefrontal cortex ‒especially its medial subdivision (mPFC)‒ is a key node within a complex neural circuitry encompassing the amygdala and the hippocampus that underlies the ability to extinguish learned defensive reactions and recall extinction memories^20, 21, 49, but see^ ^50^. Based on this line of evidence, one study^15^ administered one session of 10Hz-rTMS over the mPFC and observed enhancement of extinction learning (indexed by eyeblink startle reflexes, SCRs, and subjective ratings) which persisted until 24h during extinction recall (in eyeblink startle reflexes only). Moreover, these behavioral results were mirrored by the functional near-infrared spectroscopy (fNIRS) findings, which revealed increased mPFC activity in the stimulated group relative to the sham group. In one following work, Raij *et al.* (2018) tested the effects of an online time-locked TMS indirectly focused on the ventral mPFC (vmPFC) on the expression of threat-extinction memory. During extinction learning, the authors delivered brief 20Hz-rTMS trains over the left posterior PFC ‒a region that showed robust functional connectivity with the vmPFC. This experimental design resulted in a reduction of defensive responses during extinction recall (24h later), which was specific for the cue that had been paired with rTMS.

In our study, we tested both immediate and remote (one week) rTMS effects and we did not include extinction training before retrieval. We observed a significant decrease in defensive reactions even in the first CS trial of the test session, i.e. the first time that subjects were re-exposed to the CS after the formation of the CS-US association, and this effect was maintained in the follow-up session. Hence, since we found this outcome in absence of extinction training, it is likely that the rTMS procedure directly modulated the defensive responses activated by the implicit threat memory trace. Alternatively, the rTMS procedure over the mPFC may have inhibited the recall of the CS-US predictive association, preventing the defensive responses to be activated by the CS but leaving the responses toward the US_2_ intact. This possibility would be in line with a large body of literature on humans^see^ ^19^ which demonstrates the importance of mPFC for value-based processing, as well as with rodent studies^51^ reporting that mPFC is critical for the expression of conditioned but not innate fear.

Moreover, one core knowledge about extinction is that under certain circumstances ‒such as a simple passage of time (i.e., spontaneous recovery) or a change in surrounding context (i.e., renewal)‒ extinguished reactions triggered by the CS may reoccur, giving rise to the phenomenon known as return of fear^44, 52, 53^. To test potential renewal phenomena, which have not been investigated in the aforementioned studies^15, 16^, we opted for a context-shift amongst the learning (context A), the test (context B), and the follow-up phase (context A), and we found abolished defensive reactions in both the test and the follow-up phases. These data demonstrated that the mPFC-rTMS protocol long-term reduced threat memory expression in a different context as well as in the context in which the threatening experience had occurred, thus preventing the return of fear.

To potentiate the neural activity of the PFC, both the aforementioned studies^15, 16^ adopted high-frequency rTMS protocols ‒which are conventionally considered excitatory of proximal brain activity^54^. In our study, we adopted a low-frequency rTMS protocol ‒which is conventionally considered inhibitory^54^. Recent evidence, however, challenged this common frequency-dependent rule^55^. Indeed, resting-state functional magnetic resonance imaging (fMRI) studies demonstrated that 1Hz-rTMS protocols may also induce distal effects and enhance functional connectivity amongst the brain regions located underneath the coil and remote brain areas of the stimulated functional network^34^. Additionally, some studies^35, 36^ reported that 1Hz-rTMS procedures delivered over the PFC may paradoxically increase local brain activity.

The dorsolateral PFC is another prefrontal region that is assumed to be critically involved in threat learning^56, 57^ and the down-regulation of the cortico-meso-limbic network^58^. The vast majority of the studies targeted this site to modulate threat processing^6^. One investigation^17^ probed the effects of a state-dependent 1Hz-rTMS over bilateral dlPFC after memory reactivation to disrupt threat-memory reconsolidation. The authors observed that the stimulated groups failed to discriminate between threatening and safe stimuli (with an increase in autonomic responses to these last ones), and inferred that the perturbation of either left or right dlPFC during the reconsolidation time window was causally associated with a reduction of implicit defensive responses. A more recent study^59^ adopted the continuous theta-burst stimulation (cTBS) over the right dlPFC during the reconsolidation window and successfully decreased the defensive responses for both recent and remote threat memories. In our study, by directly comparing the effects of the 1Hz-rTMS protocol over the mPFC and the left dlPFC, we found a stronger attenuation of implicit defensive responses to the threatening cue in subjects that were stimulated over the mPFC, suggesting that targeting this brain site might represent a more promising approach for therapeutic applications.

The neural mechanisms by which rTMS over the mPFC decreases threat-conditioned responses can be manifold. Fear memories are formed and retrieved by an intricate neural network encompassing the amygdala^60^, the cerebellum^61‒63^, and sensory cortices^40, 64‒72^. Through the direct or indirect connections of the mPFC with these areas, it might be that the effects of focal manipulations of mPFC activity (i.e., stimulation or inhibition) reflect more complex and dynamic changes in the overall neural network state and/or influence the activity of some of these areas. In particular, based on the putative role of the mPFC in top-down regulating the amygdala^18^, one possible interpretation of our data is that our rTMS procedure enhanced the wide-range connectivity of mPFC and caused a downstream inhibition of the amygdala, resulting in a strong depotentiation of defensive reactions to conditioned stimuli. As an alternative to the inhibitory model of mPFC^18, 20, 73‒77^, other studies suggest that mPFC activity may underpin aversive emotions^78‒82^. According to this perspective, our rTMS protocol may have inhibited mPFC activity, thus suppressing defensive responses. Future neuroimaging studies should disentangle this issue. It should be noted that the adoption of the terms “mPFC” or “vmPFC” is not referred to brain areas delimited by clearly and univocally determined anatomical boundaries, but may rather depend on the topographical accuracy of each experimental design^19^.

Although several studies enlightened the role of the mPFC and mPFC‒hippocampus connectivity in emotion-related components of episodic memory^83^, we did not detect any rTMS-driven effect on explicit recognition memory. The observed divergence between autonomous and declarative patterns might have been due to a selective rTMS action upon the neural system supporting implicit threat processing, which has been widely dissociated from the neural system underlying explicit memory processes^39, 46‒48^. Critically, an rTMS procedure that shapes implicit overreactions to learned threats without affecting conscious knowledge of the same stimuli might represent a strategic advantage for therapeutic applications.

Since prevention of relapse is the main challenge for therapies dedicated to post-traumatic and anxiety disorders, our findings may represent an advance in this direction by providing a potential strategy to deactivate emotional overreactions and, most of all, to prevent the return of fear. Future research perspectives might consist of exploring this rTMS application over the mPFC in clinical populations displaying high levels of anxiety or suffering from anxiety disorders and PTSDs.

## Materials and Methods

### Participants

All participants (*n* = 133) were healthy volunteers (mean age: 23.62 ± 2.57, 50 males and 83 females) with no history of psychiatric disorders, neurological illnesses, cardiovascular diseases, illegal drug use, musical trainings, nor any other exclusion criteria for TMS administration^84^. During the pre-experimental screening phase, each volunteer was also administered with the *State-Trait Anxiety Inventory Form Y*^85, 86^, and who showed a score >80 in the sum of the two subscales (State + Trait anxiety) was not included in the sample (see Table 1 for all groups’ mean State-Trait Anxiety Inventory scores). Participants were then randomly assigned to each experimental condition, based on sex and age (see Table 1 for all groups’ mean age and sex distribution). We discarded seven participants because of a complete absence of skin conductance responses (SCRs) during the test session, leaving a total of 126 participants. Each participant provided written informed consent after receiving a complete description of the experimental procedures. All experimental procedures were performed in accordance with the ethical standards of the Declaration of Helsinki and were approved by the Bioethics Committee of the University of Turin.

### Auditory stimuli

Auditory stimuli were pure sine wave tones with oscillation frequencies of 800Hz (CS), 1000Hz (NS_1_), and 600Hz (NS_2_), lasting 6s with onset/offset ramps of 5ms. Tones were digitally generated using Audacity 2.1.2 software (Audacity® freeware). The unconditioned threatening stimulus (US_2_) consisted of a woman scream sample lasting 4s. All auditory stimuli were binaurally delivered through headphone speakers (Direct Sound EX29) at 50 dB intensity. All experimental scenarios were controlled by Presentation® 21.1 software (NeuroBehavioral Systems, Berkeley, CA).

### Preconditioning

This phase consisted of the presentation of 4 trials of the CS (800Hz) with an inter-trial-interval (ITI) randomly ranging between 21s and 27s. SCRs were recorded during this phase to provide a baseline response pattern to the 800Hz tone for each participant. At the end of this phase, participants were asked to confirm whether the tones were easily audible but not too loud or annoying.

### Unconditioned stimulus calibration procedure

Before starting with the calibration procedure, systolic and diastolic blood pressure was measured to prevent possible hypoarousal reactions caused by basal hypotension. The unconditioned stimulus (US) consisted of a mild electrical shock (train pulse at 50Hz lasting 200ms, with a single pulse duration of 1000µs) generated with a direct current stimulator (DS7A Constant Current Stimulator, Digitimer). Impulses were delivered through a bar stimulating electrode connected by a Velcro strap on the upper surface of the dominant hand’s index finger. The electrical stimulation intensity was individually calibrated through a staircase procedure^39, 45, 87^, starting with a low current near the perceptible tactile threshold (∼0.5 mA). Participants were asked to rate the painfulness of each train pulse on a scale ranging from 0 (not painful at all), 1 (pain threshold) to 10 (highly painful if protracted in time). At the end of the procedure, the US amplitude was then set at the current level (mA) corresponding to the mean rating of ‘7’ on the subjective analog scale.

### Conditioning

After a 1-min resting period, participants underwent a single-cue auditory threat conditioning, which consisted of the presentation of 15 trials of the conditioned stimulus (CS, 800Hz), with an ITI randomly ranging between 21s and 27s. The CS co-terminated with the US 12 times (80% reinforcement rate). Subjects were not informed about any possible CS-US contingency. To validate the threat learning experience, immediately following this phase subjects rated the painfulness of the US using the same analog scale as in the preconditioning calibration procedure (see Table 1 for all groups’ US current intensity and US analog ratings).

### Transcranial Magnetic Stimulation

Transcranial Magnetic Stimulation was performed with a Magstim Rapid^2^ Stimulator (Magstim Co., Whitland, Dyfed, UK). A 70-mm figure-of-eight coil was positioned over the subject’s M1 cortical area at the optimum scalp position to elicit a contraction of the contralateral abductor *pollicis brevis* muscle (APB). Resting motor threshold (rMT) was defined as the minimum stimulation intensity that induced a visible finger movement in at least 5 out of 10 single pulses over the right hand area of the left primary motor cortex^15, 88^. After having determined each individual’s rMT, we applied a single train of 1Hz-rTMS for a total duration of 10min (600 pulses) to the target area. The rTMS intensity was set at 80% of the rMT for subjects whose rMT was ≤ 50% of the machine’s maximum deliverable power (e.g., the intensity corresponded to 40% of the maximum power when the rMT was equal to 50% of the same parameter). For subjects with an rMT > 50%, the stimulation intensity was always set to a ceiling corresponding to 40% of the machine’s maximum deliverable power (see Table 1 for each group’s mean rMT and mean stimulation intensity). During the rTMS procedure participants were seated in a comfortable recliner that we adjusted to allow their upper body to be in a sloped position, thus ensuring an optimal positioning of the coil.

To target the medial portion of the prefrontal cortex (BA 10; mPFC and mPFC-E groups), the coil was centered over Fpz (10% of nasion-inion distance) according to the international 10‒20 electroencephalogram (EEG) system^89^ (Figure 1). This placement should ‒with an rTMS reach of 1.5 to 2 cm beneath the scalp^90, 91^‒ ensure the targeting of the mPFC as in previous studies^15, 32, 33^ and avoid the targeting of the dorsomedial prefrontal cortex (dmPFC), which would have been localizable with a scalp-based heuristic approach of 25.84% nasion-inion distance^92^. In the case of left occipital cortex stimulation (OC group), the coil was positioned over O1 using the 10–20 EEG system (BA 18/19), which functionally corresponds to associative visual cortices V3, V4, and V5^93, 94^ (Figure 1). For the stimulation of the left dorsolateral prefrontal cortex (dlPFC group), the coil was placed over F3 using the 10–20 EEG system (BA 8/9)^17, 95^ (Figure 4A). For sham stimulation (sham and sham-E groups), the coil was centered over Fpz and positioned perpendicular to the scalp surface, so that no effective stimulation reached the brain during the procedure but allowed subjects to feel a comparable coil-scalp contact and hear the same noise as in real stimulation (Figure 1).

All participants were blinded to their experimental condition (i.e., active or sham), and were not informed about the potential cognitive or emotional effects of the stimulation.

### Implicit recognition test

After a 1-min resting period participants underwent this task, which consisted of the presentation of 12 auditory stimuli in a completely random sequence: 4 × CS, 4 × NS_1_, 4 × NS_2_, with an ITI whose duration randomly ranged between 21s and 27s. SCRs were recorded throughout this phase, and the stimulating electrode was kept attached to create the expectation to receive the US^37^. Differently from other paradigms^42, 96‒98^, here no shocks were delivered to avoid any reacquisition effect^39, 45^.

### Implicit unconditioned threatening test

This task was designed to elicit an unconditioned electrodermal response and consisted of the presentation of 4 trials of a woman scream sample lasting 4s, with an ITI randomly ranging between 21s and 27s. SCRs were recorded throughout this phase.

### Two-alternative forced-choice (2AFC) explicit recognition test

This procedure involves the presentation of two stimuli on each trial and the subject chooses the one that was previously encoded (i.e. the first or the second one). As in our previous works^39, 45^, a 2AFC design was preferred over a new-old paradigm, which involves one single stimulus on each trial and the subject judges whether the stimulus has been previously encoded (old), or whether it is new. Our choice was motivated by the evidence that a 2AFC task improves recognition performance and discourages response biases such as the familiarity-based decision bias, namely the heuristic to endorse novel cues as ‘old’ when their familiarity is high^99^.

The task consisted of the presentation of 16 tone-pairs, each composed of the CS (800Hz) and one of the two NSs (NS_1_, 1000Hz or NS_2_, 600Hz) in a completely random sequence: 4 × CS vs NS_1_, 4 × NS_1_ vs CS, 4 × CS vs NS_2_, 4 × NS_2_ vs CS. On each trial, the two stimuli were presented with an intra-trial-interval of 1000ms. After each pair offset, an ITI randomly ranging between 21s and 27s occurred. Participants were explained that in each couple of sounds there was a tone that they had heard on the first session (one week before or, in the case of the follow-up session, two weeks before) and a new tone. Participants were then instructed to recognize and verbally refer which one (the first or the second) was the tone heard in the first session, paired with the US-shock (CS). Participants were further asked to verbally provide a confidence rating about each response, on a scale from 0 (completely unsure) to 10 (completely sure). No feedback was supplied. As in the implicit task, the stimulating electrode was kept attached, but no shock was delivered.

### Two-alternative forced-choice (2AFC) perceptual discrimination test

The task consisted of the presentation of 7 pairs of auditory stimuli (i.e. CS vs NS_1_, NS_1_ vs CS, CS vs NS_2_, NS_2_ vs CS, CS vs CS, NS_1_ vs NS_1_, NS_2_ vs NS_2_) with a 1000-ms intra-pair-interval in a completely random sequence (ITI randomly ranging between 21s and 27s). For each pair, subjects were asked to refer whether the two tones were “the same tone or different tones”, and to provide a confidence rating on an analog scale from 0 (completely unsure) to 10 (completely sure). No feedback was supplied.

### Psychophysiological recording and analysis

Event-related skin conductance responses (SCRs) were used as an implicit index of defensive responses. To record the autonomic signal, two Ag-AgCl non-polarizable electrodes filled with isotonic paste were attached to the index and middle fingers of the non-dominant hand by Velcro straps. The transducers were connected to the GSR100C module of the BIOPAC MP-150 system (BIOPAC Systems, Goleta, CA) and signals were recorded at a channel sampling rate of 1000 Hz. SCR waveforms were analyzed offline using AcqKnowledge 4.1 software (BIOPAC Systems, Goleta, CA), and were performed blindly to the subject’s experimental condition and the randomized sequence of stimuli. Each SCR was evaluated as event-related if the trough-to-peak deflection occurred 1–6 s (for the CS and the NSs) or 1–4 s (for the US_2_) after the stimulus onset, the duration was comprised between 0.5 and 5.0 s, and the amplitude was greater than 0.02 micro siemens (μS). Responses that did not fit these criteria were scored zero. Raw SCR data were square-root transformed to normalize the distributions^100^. To account for inter-individual variability, these normalized values were then scaled according to each participant’s unconditioned response by dividing each response by the mean square-root transformed unconditioned stimulus response^101, 102^. To obtain a quantification of each participant’s post-conditioning response level, we finally calculated the delta score between these scaled values and each subject’s mean square-root transformed response to the 800Hz-tone (CS) prior to the conditioning (i.e. preconditioning phase).

### Statistical analyses

We computed the appropriate sample size based on a power analysis performed through G*Power 3.1.9.2. For the main statistics, i.e. one-way ANOVA with three groups, with the following input parameters: α equal to 0.05, power (1-β) equal to 0.80, and a hypothesized effect size (η_p_^2^) equal to 0.14, the estimated sample size resulted in *n* = 21 per experimental group, which was larger than those adopted in previous TMS studies^17, 59^.

Since all variables passed the D’Agostino-Pearson omnibus normality test, parametric statistics were adopted in each experiment. To test the between-groups differences in post-conditioning US ratings, preconditioning SCRs levels, and SCRs to the US during conditioning, we performed one-way ANOVA models and Student’s unpaired *t* tests. To test the potential between-groups (sham, OC, and mPFC) differences in the implicit reactions to the CS, to the NS_1_, to the NS_2_, and the US_2_ during the test session, and the implicit reactions to the CS and the US_2_ during the follow-up session, we computed one-way ANOVAs with Bonferroni-adjusted *post hoc* comparisons. To test potential differences in the time course of CS-related responses, we performed 3 × 4 mixed ANOVAs with Group (sham, OC, and mPFC) as between-subjects variable and Trial (1–4) as within-subjects variable. Bonferroni adjustment was applied for simple main effects analyses. To compare within-groups responses to the CS and NSs in each group, we performed repeated measures ANOVAs.

To test the between-groups differences in the explicit recognition and respective confidence ratings, as well as in the perceptual discrimination and respective confidence ratings during the test and the follow-up sessions, we performed Student’s unpaired *t* tests. To test whether explicit recognition levels were significantly higher than the 50% chance level for each condition during the test and the follow-up sessions, we calculated Student’s one sample *t* tests against 0.50.

To test the between-groups (dlPFC vs mPFC) differences in autonomic reactions to the CS and the NSs during the test, and to the CS during the follow-up, we adopted Student’s unpaired *t* tests. To test potential differences in the time course of CS-related responses, we performed 2 × 4 mixed ANOVAs with Group (dlPFC and mPFC) as between-subjects variable and Trial (1–4) as within-subjects variable. For each ANOVA we assessed the Sphericity assumption through Mauchly’s Test. Where it was violated, we applied the Greenhouse-Geisser correction accordingly.

The null hypothesis was rejected at *P* < 0.05 significance level. All statistical analyses were performed using SPSS Statistics 22 (IBM) and Prism 9 (GraphPad).

## Acknowledgements

We thank all the subjects for their participation in this study. We also thank Melania Lattuada, Veronica Cintori, Ester Fusaro, Alessia Di Blasi, and Adriana Monno for their help in the experimental procedures and for the continuous support and advice. This work was supported by the “Compagnia di San Paolo, Progetto d’Ateneo”, University of Turin 2017 (CSTO167503), by the Fondazione Giovanni Goria and Fondazione CRT (Talenti della Società Civile, ed. 2018), by the Grant “Progetti di ricerca di Rilevante Interesse Nazionale (PRIN)” 2017 (Project n. 20178NNRCR_002) from the Italian Ministry of University and Research (MIUR), Fondazione Cariverona 2018, and Fondazione CRT 2021.

## Supplementary Information

### Supplementary Figures

**Supplementary Figure 1.**
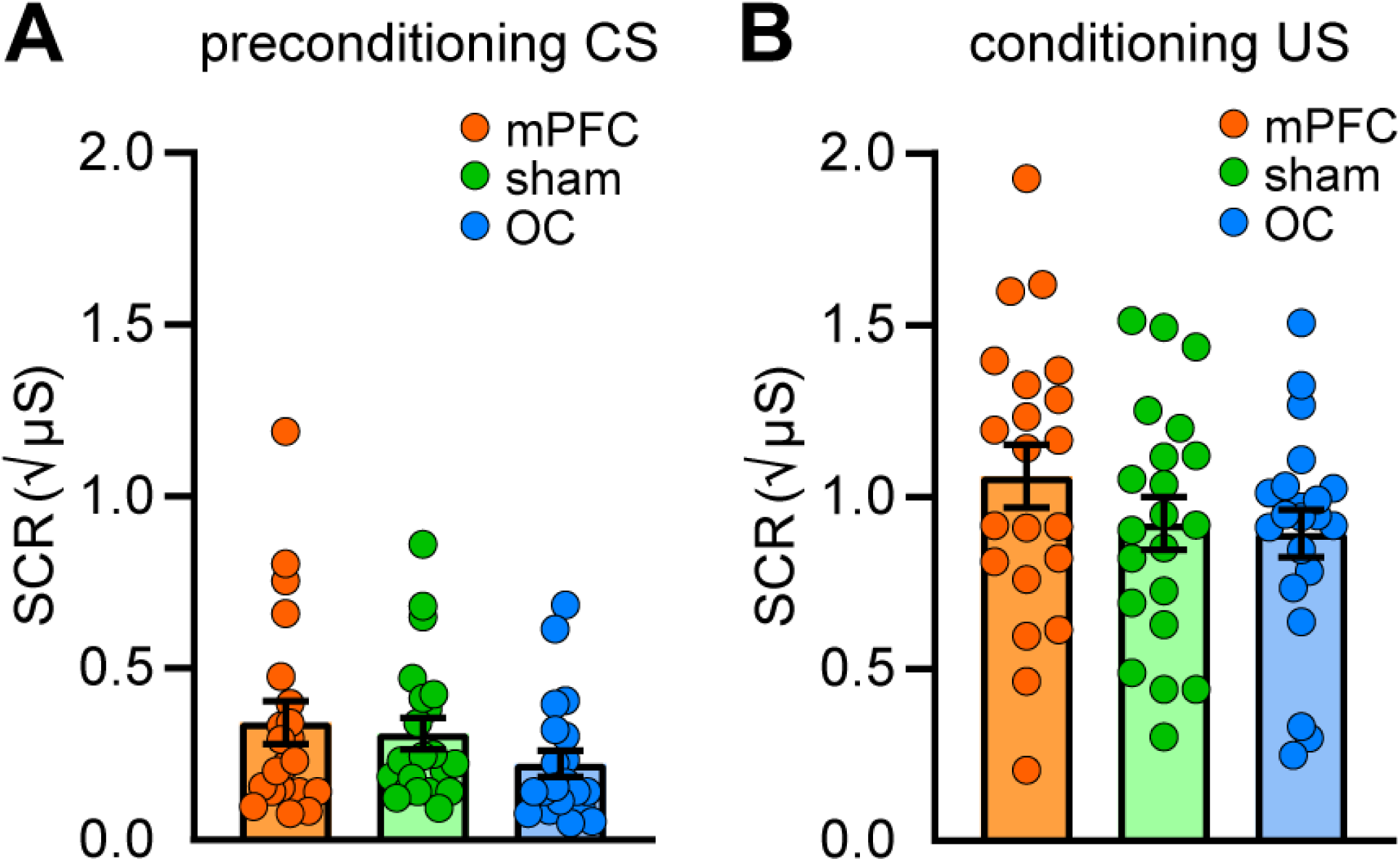
Implicit reactions of mPFC-stimulated, OC-stimulated, and sham groups during preconditioning and conditioning. (**A**) Dot plot representing the mean SCRs elicited by the CS during the preconditioning phase in the three different conditions (mPFC, *n* = 21; sham, *n* = 21; OC, *n* = 21). Implicit reactions were similar among conditions. (**B**) Dot plot representing the mean SCRs elicited by the US during the conditioning phase in the three different conditions. Implicit reactions were similar among conditions. All data are mean and SEM. One-way ANOVA (A, B).

**Supplementary Figure 2.**
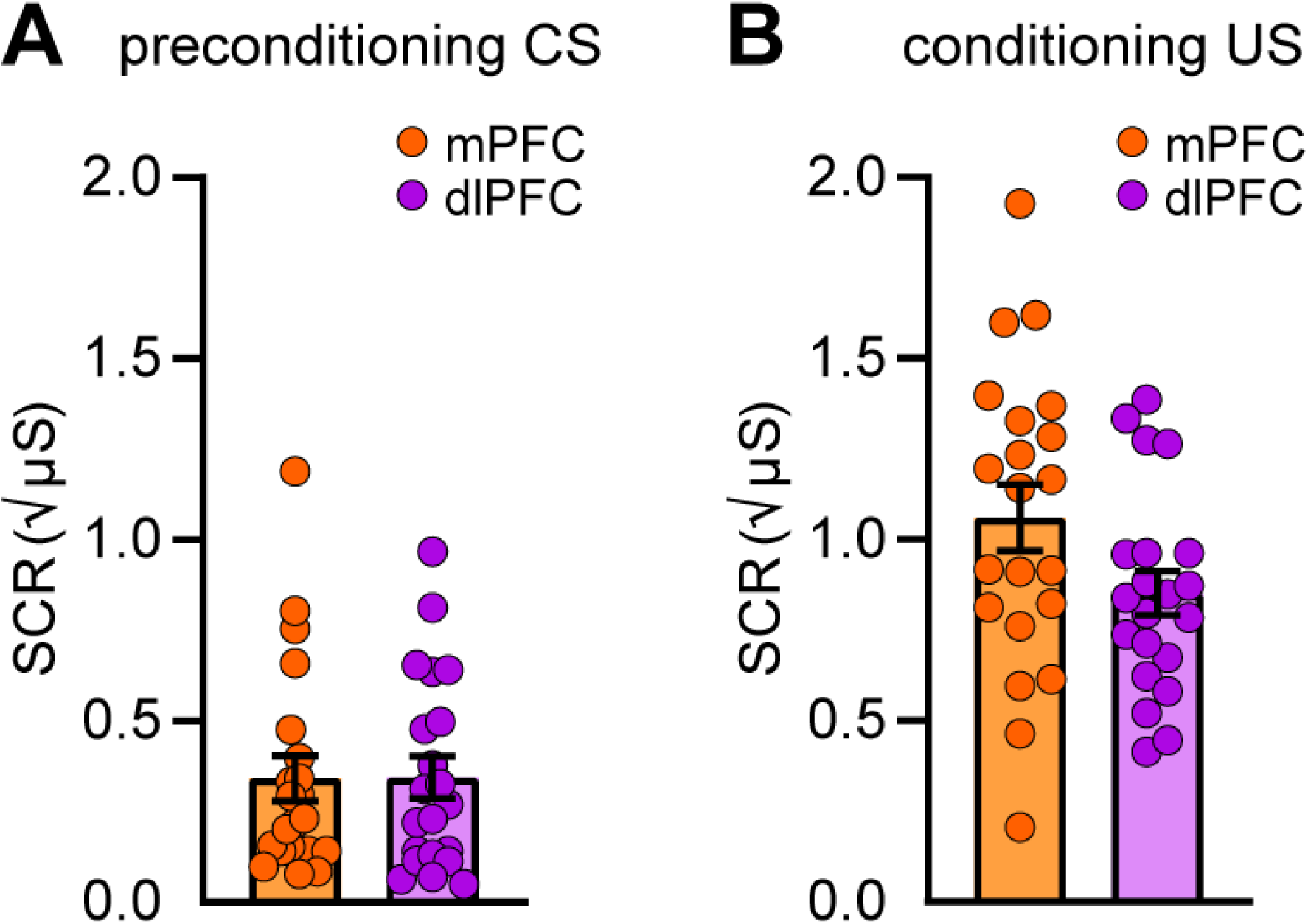
Implicit reactions of mPFC-stimulated and dlPFC-stimulated groups during preconditioning and conditioning. (**A**) Dot plot representing the mean SCRs elicited by the CS during the preconditioning phase in the same mPFC group as in Supplementary Figure 1 and the dlPFC group (*n* = 21). Implicit reactions were similar between conditions. (**B**) Dot plot representing the mean SCRs elicited by the US during the conditioning phase in the two different conditions. Implicit reactions were similar between conditions. All data are mean and SEM. Student’s unpaired *t* test (A, B).

